# Exceptional origin activation revealed by comparative analysis in two laboratory yeast strains

**DOI:** 10.1101/2021.02.16.431410

**Authors:** Jie Peng, Ishita Joshi, Gina Alvino, Elizabeth Kwan, Wenyi Feng

**Affiliations:** Department of Biochemistry and Molecular Biology, SUNY Upstate Medical University, 750 East Adams Street, Syracuse, NY 13210; Department of Genome Sciences, University of Washington, Box 355065, 1705 NE Pacific Street, Seattle, WA 98195

**Keywords:** Origin of replication, Replication checkpoint, *RAD53*, Autonomously Replicating Sequence (ARS), ARS-consensus sequence (ACS), Single-stranded DNA (ssDNA), Single nucleotide polymorphism (SNP), A364a, W303, *Saccharomyces cerevisiae*

## Abstract

We performed a comparative analysis of replication origin activation by genome-wide single-stranded DNA mapping in two common laboratory strains of *Saccharomyces cerevisiae* challenged by hydroxyurea (HU), an inhibitor of the ribonucleotide reductase. By doing so we gained understanding of the impact on origin activation by three factors: replication checkpoint control, DNA sequence polymorphisms, and relative positioning of origin and transcription unit. Our analysis recapitulated the previous finding that the majority of origins are subject to checkpoint control by the Rad53 kinase when cells were treated with HU. In addition, origins either subject to Rad53 checkpoint control or impervious to it are largely concordant between the two strains. However, these two strains also produced different dynamics of origin activation. First, the W303-*RAD53* cells showed a significant reduction of fork progression than A364a-*RAD53* cells. This phenotype was accompanied by an elevated level of Rad53 phosphorylation in W303-*RAD53* cells. Second, W303-*rad53K227A* checkpoint-deficient cells activated a greater number of origins accompanied by global reduction of ssDNA across all origins compared to A364a-*rad53K227A* cells; and this is correlated with lower expression level of the mutant protein in W303 than in A364a. We also show that sequence polymorphism in the consensus motifs of the replication origins plays a minor role in determining origin usage. Remarkably, eight strain-specific origins lack the canonical 11-bp consensus motif for autonomously replicating sequences in either strain background. Finally, we identified a new class of origins that are only active in checkpoint-proficient cells, which we named “Rad53-dependent origins”. The only discernible feature of these origins is that they tend to overlap with an open reading frame, suggesting previously unexplored connection between transcription and origin activation. Our study presents a comprehensive list of origin usage in two diverse yeast genetic backgrounds, fine-tunes the different categories of origins with respect to checkpoint control, and provokes further exploration of the interplay between origin activation and transcription.

**Author Summary:** Comparative analysis of origins of replication in two laboratory yeast strains reveals new insights into origin activation, regulation and dependency on the Rad53 checkpoint kinase.

## Introduction

Eukaryotic genomes are reliant on multiple initiation sites, called origins of replication, on each chromosome for complete duplication of the DNA in every cell cycle. Much of our knowledge concerning the location and usage of replication origins comes from studies using the budding yeast, *Saccharomyces cerevisiae*, where origins were initially identified in a genetic assay as autonomously replicating sequences (ARS) capable of supporting plasmid replication [1]. Within each ARS, an 11-basepair ARS-consensus sequence (ACS), (T/A)TTTAT(A/G)TTT(T/A), has been found to be essential for origin function [2, 3]. The ACS serves as the binding site for the hexameric origin recognition complex (ORC) [4-7]. The ORC-bound sites are then joined by Cdc6 and Cdt1, which further recruit the hexameric MCM2-7 DNA helicase to form a pre-replicative complex (pre-RC) [8]. In the *S. cerevisiae* genome more than 12,000 ACSs have been found; however, only approximately 400 origins are used during a single cell cycle, suggesting cis-acting sequences other than the ACS function in specifying or influencing origin usage [9, 10]. For instance, a 19-bp sequence in the *URA3* locus has been found to advance the time of origin activation through an unknown mechanism [11-13]. Genetic variation in diverse laboratory strains can manifest in distinct physiological properties [14-17]. Thus, *a priori* sequence polymorphism among different laboratory strains is expected to exert a direct impact on origin usage.

Studies using a replication timing assay based on the Meselson-Stahl density transfer method revealed a hierarchy of origin activation, which follows a temporal order in S phase [18-20]. This hierarchy is more starkly manifested in cells undergoing S phase in the presence of the ribonucleotide reductase inhibitor, hydroxyurea (HU), where initiation is only observed at origins that normally fire in roughly the first half of an unchallenged S phase but not at origins that would normally fire later in the unchallenged S phase [21-24]. This hierarchy of origin activation is disrupted in cells carrying mutations in DNA replication checkpoint genes, such as *mec1* and *rad53*, in conjunction with replication stress by HU or other DNA damage-inducing drugs [21-27]. Specifically, those origins that tend to be activated later in S phase in checkpoint-proficient cells are prematurely activated in the checkpoint mutants [21, 22, 28]. Consequently, virtually all origins fire in HU in checkpoint mutants, in contrast to wild type cells where only a subset of origins fire. This phenomenon permitted the classification of origins into two groups, “Rad53-unchecked” and “Rad53-checked” origins, based on their activation status (positive and negative, respectively) in a kinase-dead *rad53K227A* mutant versus a *RAD53* background, in the presence of HU [22]. While this mechanism of checkpoint control of origin activation appears to generally hold true among different strains, whether a given origin is subject to checkpoint control equally across diverse genetic background has not been assessed in a systematic manner.

In *S. cerevisiae,* origins predominantly reside in intergenic regions [29]. This feature is presumably the evolutionary result of minimizing replication-transcription conflict, which has precedents in both prokaryotic and eukaryotic genomes. For instance, the *E. coli* genes are However, a study that systematically mapped Mcm2-7 binding sites in both mitotic and meiotic cells showed that 106 of 393 origins that contained Mcm2-7 binding sites were intragenic, *i.e.*, overlapping with the ORF or the promoter of a gene [30]. Twenty of these intragenic origins (19%) showed meiosis- or mitosis-specific Mcm2-7 loading. This differential origin activity is primarily due to gene expression during meiosis precluding origin activation and resulting in mitosis-specificity, and vice versa, underscoring transcription-replication incompatibility [30]. Nevertheless, 86 intragenic origins (81%) were apparently active regardless of meiotic or mitotic growth. How these intragenic origin activities are coordinated with transcription so as not to be obliterated by transcription is a fascinating problem. On the other hand, a recent study has shown that the ACS element(s) within the replication origin can pause or terminate RNA PolII-mediated pervasive transcription, suggesting that intragenic origins might assume a more proactive role than previously thought with regards to coordination with transcription [31]. The genetic determinants for the regulation of intragenic origins are not understood.

Driven by all the questions above we set out to systematically map and characterize origin usage in two commonly used strains, A364a and W303, using a previously described microarray-based genome-wide labeling of replication fork-associated single-stranded DNA (ssDNA) [22, 32]. Because the presence of HU restricts the pool of dNTPs and thereby limits replication fork progression, the genomic locations of S-phase specific ssDNA accumulation in HU mark the locations of active origins. We introduced a kinase-dead and checkpoint-deficient K227A mutation in the *RAD53* gene in each of the two strains and compared origin usage in wild type and the *rad53* mutant cells. We also integrated whole genome sequencing data to identify single nucleotide polymorphisms (SNPs) in the two strains. Our study demonstrates that by and large the replication checkpoint exerts control over origin activation similarly in the two strains. However, we did find strain-specific origin usage and the potential contributing single nucleotide polymorphisms (SNPs), thus enabling future studies to identify the regulatory mechanisms for origin usage. Finally, we identified a new class of origins that are only active in cells with an intact checkpoint. These origins tend to reside in regions overlapping open reading frames (ORFs). However, the lack of origin activation in the *rad53* mutant cannot be explained by increased transcription. Therefore, we suggest that origin activation at these specific locations requires an intact Rad53 kinase at an undefined step.

## Results

### Differential dynamics of origin activation in the A364a and W303 background

We surveyed origin activity in cells containing wild type *RAD53* or a *rad53K227A* kinase-dead mutation in both genetic backgrounds, A364a and W303, using a previously described method of mapping ssDNA gaps at the replication fork stalled by HU [22, 32]. Briefly, yeast cells were embedded in agarose plugs and then spheroplasted to reveal the chromosomal DNA while protecting the single-stranded gaps therein. The ssDNA would then serve as the template for random-primed DNA synthesis without denaturation to separate the duplex DNA. The DNA from an S phase sample and a G1 control sample were differentially labeled with Cy3- and Cy5-dNTP, respectively, and then co-hybridized to a microarray to reveal the locations of S-phase specific ssDNA (Fig. 1A). Two biological replicates were performed for each sample. In all cases the biological replicates exhibited high concordance (R>=0.97), underscoring the reproducibility of the ssDNA measurements (Fig. 1B). Chromosomal plots of ssDNA profiles were nearly identical between replicates, revealing ssDNA peaks at discrete locations of the chromosomes (Fig. 1C). See all chromosome plots for Exp 1 in Fig. S1.

**Figure 1.**
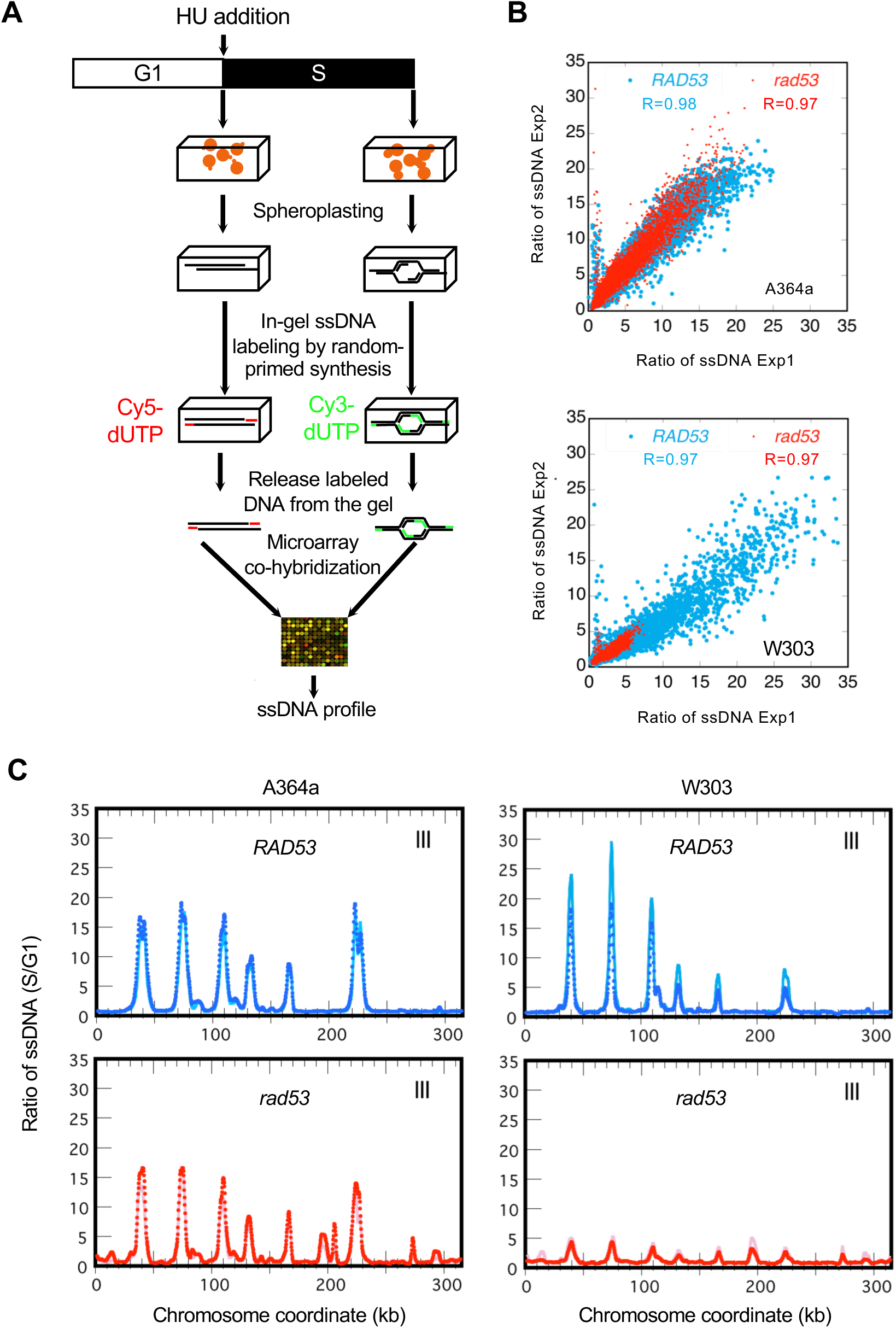
Genome-wide ssDNA mapping in two laboratory yeast strains. **(A)** Schematic experimental procedures of ssDNA mapping by microarray. (**B**) Reproducibility of ssDNA ratios (S/G1) from two independent experiments. Pearson correlation co-efficient values are shown. **(C)** Normalized ssDNA profiles of ChrIII (ssDNA ratios of S/G1 plotted against chromosome coordinates in kb) for *RAD53* and *rad53-K227A* cells in the A364a (**left**) and the W303 (**right**) background from one of the two replicate experiments (Exp 1).

Next, we developed a pipeline to systematically query origin firing status based on ssDNA levels at each of 626 “confirmed” and “likely” origins queried at OriDB (http://cerevisiae.oridb.org/). The two basic components of the pipeline were 1) ssDNA peak calling by local maxima identification and 2) assigning the ssDNA signals to a particular origin. A typical origin in wild type yeast sends off a pair of replication forks in opposite directions, resulting in a ssDNA profile of two adjacent peaks flanking the center of an origin. However, the same ssDNA profile can be mistaken for two adjacent origins each producing a single ssDNA peak. Therefore, a major challenge was to accurately differentiate these two possibilities and assign the ssDNA signals to the correct origin(s). To do so we employed three rigorously chosen and stringent parameters (Materials and Methods). First, only those origins with a ssDNA peak amplitude greater than three standard deviations above background in both replicate experiments were considered. Second, we uniformly defined origin size to be 4 kb to enable optimal ssDNA peak-origin association in an unbiased fashion. Third, we determined a minimal distance of 1.75 kb between two adjacent ssDNA peaks to be accepted as two separate origins. As detailed in Materials and Methods, our algorithm would incur an estimated 4% false negative rate due to the chosen 1.75 kb minimal inter-peak distance. Additionally, origins with moderate level of ssDNA might also elude detection, as in the case of *ARS813* (see below). Overall, we identified 362 and 405 consensus origins (detected in both replicate experiments) for A364a and W303, respectively (Fig. 2A and File S1). Altogether there were 419 unique origins of which the majority (348) were shared by A364a and W303 and the rest were activated in a strain-specific manner. Each of the 419 origins was then classified into one of three categories based on the comparison between *RAD53* and the *rad53K227A* mutant (see below).

**Figure 2.**
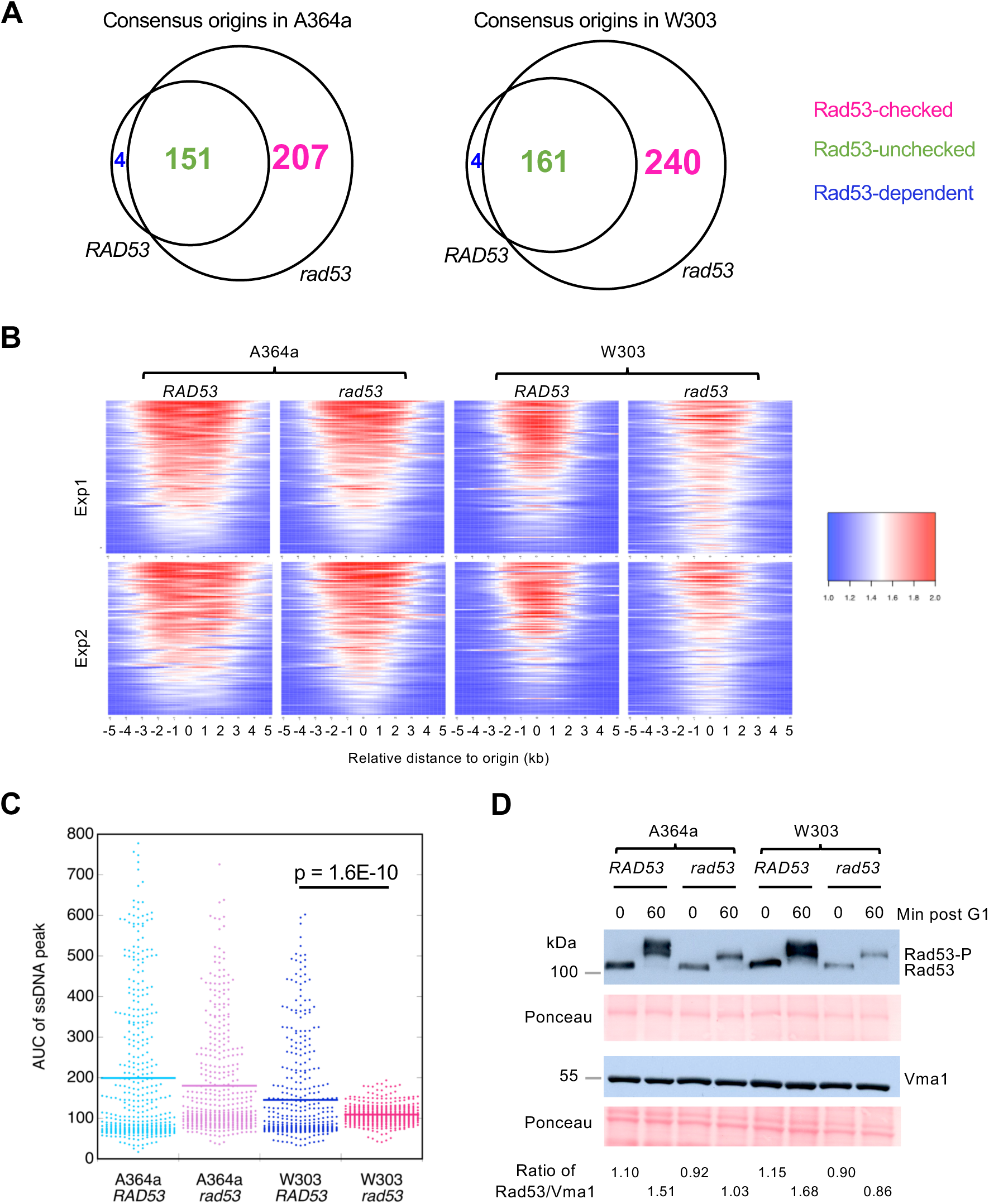
Comparative analysis of origins of replication in the A364a and W303 background. **(A)** Consensus origins identified by both biological replicate experiments in the A364a (left) and W303 (right) background, respectively, were divided into three classes of origins, “Rad53-unchecked”, “Rad53-checked”, and “Rad53-dependent”, based on the comparison between *RAD53* and *rad53* cells. See text for detail. (**B)** Heatmaps of the normalized ssDNA ratios (compressed to a scale of 1 to 2) from the 419 unique origins from (A) in each of the four strains. **(C)** Quantification of average AUC values for the uncompressed ssDNA signals at the 419 origins in each strain from two experiments. **(D)** Western blots of Rad53 levels in cells at 0 or 60 min post release from G1 arrest into S phase in the presence of 200 mM HU, respectively. A cellular protein Vma1 was used as loading control. Overall protein levels were also controlled by Ponceau staining. The Rad53 level is quantified as the ratio of Rad53/Vma1.

We first compared the extent of ssDNA progression over all 419 origins in each strain by visualizing them on a heatmap. The results showed that the extent of ssDNA formation in both *rad53* mutants in the A364a and W303 background was reduced compared to their *RAD53* counterparts (Fig. 2B). This is consistent with the notion that checkpoint failure causes replication forks to collapse shortly after initiation from the origin. Additionally, both the *RAD53* and mutant versions of the W303 background showed reduction of ssDNA compared to their A364a counterparts (Fig. 2B). Similar conclusions were drawn from generating aggregated ssDNA profiles across a 20 kb window centering on all 419 origins in each experiment (Fig. S1A) followed by calculating area under the curve (AUC) of the ssDNA peaks and averaging two replicate experiments for each sample (Fig. 2C). Additionally, *rad53* mutant exhibited global reduction of ssDNA at origins compared to wild-type cells specifically in the W303, and not the A364a background (Fig. 2C). We also calculated the average fork distance based on the width of the ssDNA peaks for each sample (Fig. S1B). Fork distance was significantly reduced in W303-*RAD53* cells than in A364a-*RAD53* cells (Fig. S1A&B). These results were not due to a difference in synchrony of the cell culture as cell cycle analysis using budding index (Fig. S2A) or flow cytometry (Fig. S2B) both confirmed synchronous entry into S phase. We then asked if the reduced fork progression in W303 is attributable to a relatively more active checkpoint by analyzing the Rad53 phosphorylation level. Indeed, in the checkpoint-proficient cells phospho-Rad53 (Rad53-P) after 60 min in HU was more abundant in W303 than A364a (Fig. 2D). The K227A mutation significantly reduced the levels of both Rad53K227A and Rad53K227A-P regardless of strain background (Fig. 2D).

### Origins with different checkpoint status in A364a and W303

We next defined the unique status for each of the 419 origins based on its behavior in the four samples, *i.e.*, in *RAD53* and *rad53* cells of either A364a or W303 background. This resulted in the classification of the 419 origins into 15 categories (File S1). First, we focused on the “Rad53-checked origins” and “Rad53-unchecked origins” (Fig. 3A), previously defined as those origins only activated in *rad53* cells, hence subject to Rad53 checkpoint control, and those shared by *RAD53* and *rad53* cells, hence oblivious to Rad53 checkpoint control, respectively [22]. One hundred and forty-seven of 151 Rad53-unchecked origins in A364a (97%) were also identified as Rad53-unchecked origins in W303 (category 1 in Fig. 3A). Similarly, 91% (185 of 207) of Rad53-checked origins in A364a were also found as Rad53-checked origin in W303 (category 2 in Fig. 3A). Therefore, the vast majority of origins share the same checkpoint status despite differences in genetic background. Only twelve origins showed opposite classifications (sum of categories 3 and 4 in Fig. 3A), which we referred to as “differential checkpoint status” origins.

**Figure 3.**
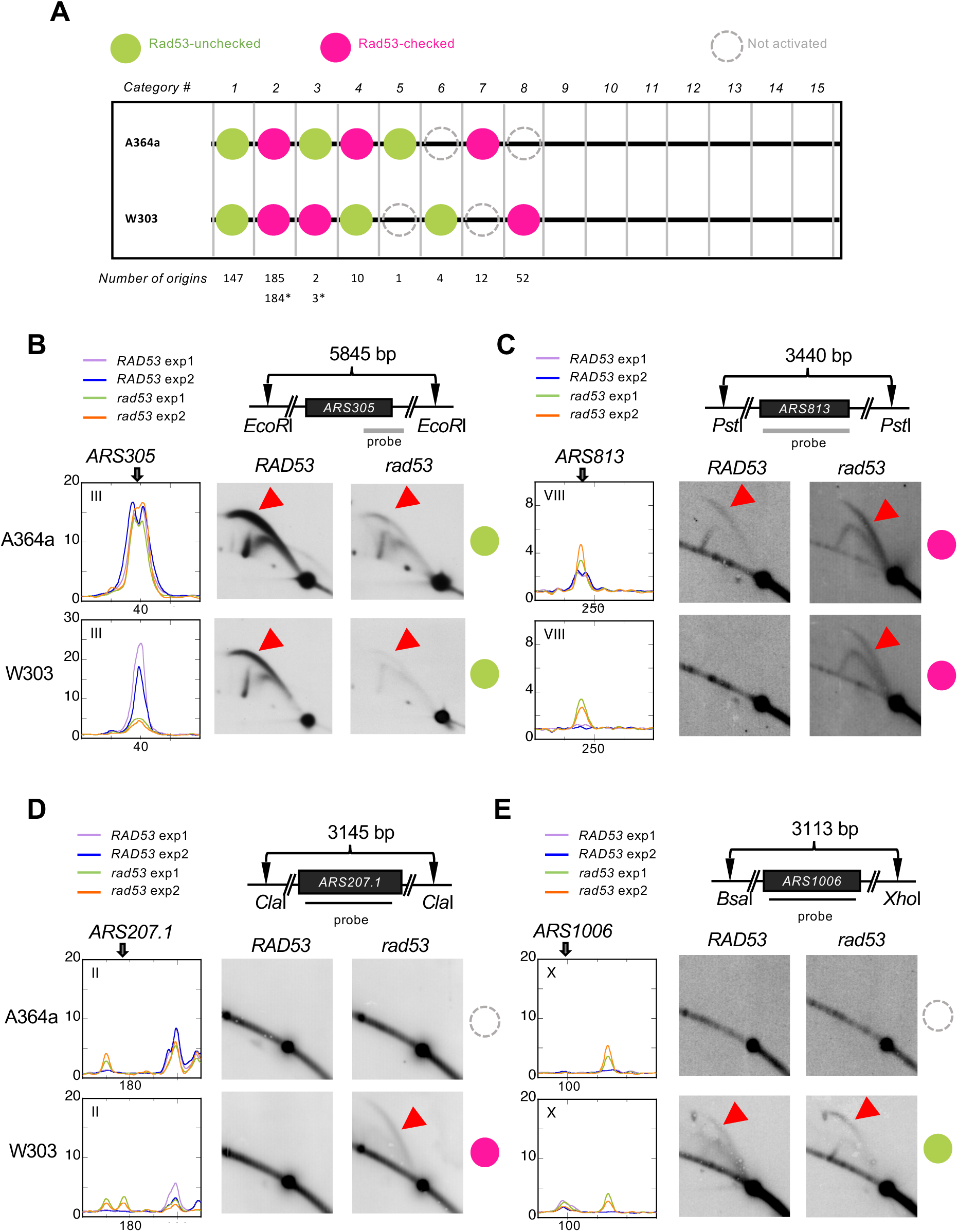
Validation of Rad53-checked/unchecked origin firing status using 2-D agarose gel electrophoresis. (**A**) Comparison of origin usage in the two strains leads to further classification of active origins into 15 categories. The number of origins in each category is listed below the diagram. The numbers with asterisks are the adjusted number after manual inspection. See text for detail. (**B-E**) DNA fragments containing select origins for analysis (*ARS305, ARS1006, ARS207.1,* and *ARS813*) were produced by restriction digestion with the indicated restriction enzymes. DNA probes are shown as black bars. Origin position is as indicated on the ssDNA profiles from two biological replicate experiments. “Bubble arcs” indicative of origin activation are indicated as red arrowheads. The origin classes were indicated using the key described in (**A**).

We verified these results by 2-D gel analysis, using the same conditions employed for the ssDNA mapping experiments, *i.e.*, harvesting cells synchronously released from G1 arrest into S phase in the presence of HU for 60 min. We did not use asynchronous cell populations to maximize the chance of detecting strain-specific differences as a result of HU-induced replication stall. We first validated *ARS305*, a Rad53-unchecked origin in both W303 and A364a. As expected, bubble arcs were detected in all samples, with or without an intact checkpoint (Fig. 3B). Notably, the bubble arc signal in *rad53* cells in W303 was significantly reduced compared to other samples, confirming the observed global reduction of origin activation based on ssDNA mapping. We also analyzed *ARS813*, which was classified as a Rad53-checked origin in both A364a and W303 and yet showed a clear ssDNA signal in *RAD53* cells in A364a (Fig. 3C). Indeed, bubble arcs were detected in this the A364a-*RAD53* cells (Fig. 3C). Thus, *ARS813* was mis-classified as category 2 by the algorithm due to the moderate level of ssDNA (less than three standard deviations above background), and should fall in category 3. This brought the number of “differential checkpoint status” origins to 13 (sum of categories 3, with asterisk, and 4 in Fig. 3A).

### Strain-specific usage of origins

We identified 71 (14 in A364a and 57 in W303) origins that were only activated in one of the two strains (sum of categories 5-8 in Fig. 3A and categories 12 and 15 in Fig. 5A, discussed later). We termed them “strain-specific origins” and also confirmed them by 2-D gel analysis. For instance, a bubble arc was detected at *ARS207.1* only in *rad53* cells in the W303 background, making it a W303-specific and Rad53-checked origin (Fig. 3D). Similarly, a bubble arc was detected at *ARS1006* in both Rad53 and *rad53* cells in W303, but not in A364a, making *ARS1006* a W303-specific and Rad53-unchecked origin (Fig. 3E).

**Figure 4.**
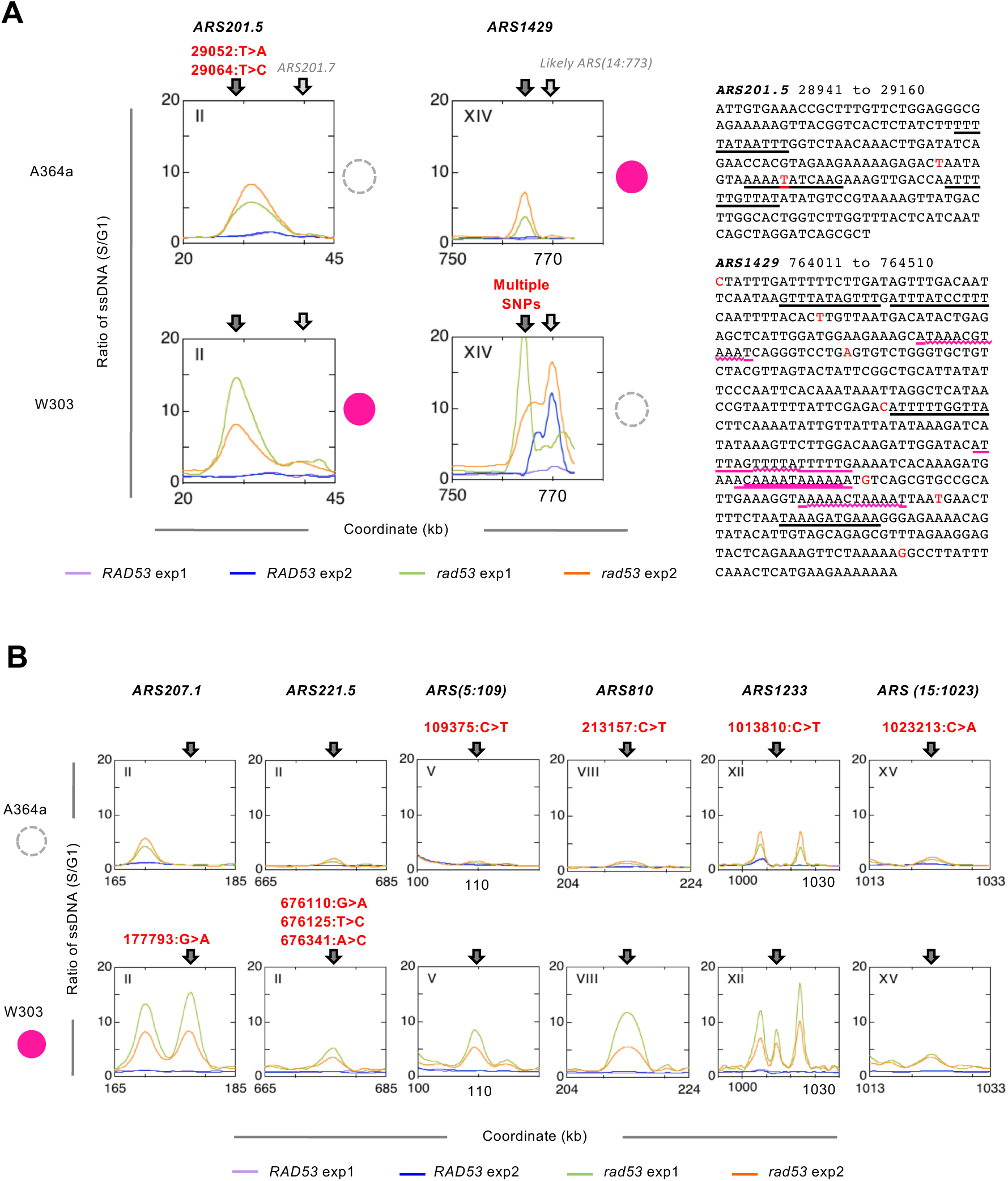
Examples of strain-specific origins with polymorphisms. The corresponding origin status is color coded same as in Fig. 3. **(A)** Two origins with polymorphisms in A364a or W303 background resulting in a shift of initiation sites from the canonical origins. The SNPs (red font) are indicated in the ARS regions based on OriDB curation on the right. The near-matches of ACS elements are underlined in black, with overlapping elements underlined in magenta. **(B)** Six origins that show binary pattern of activation in one background but not the other, without shifting initiation sites.

**Figure 5.**
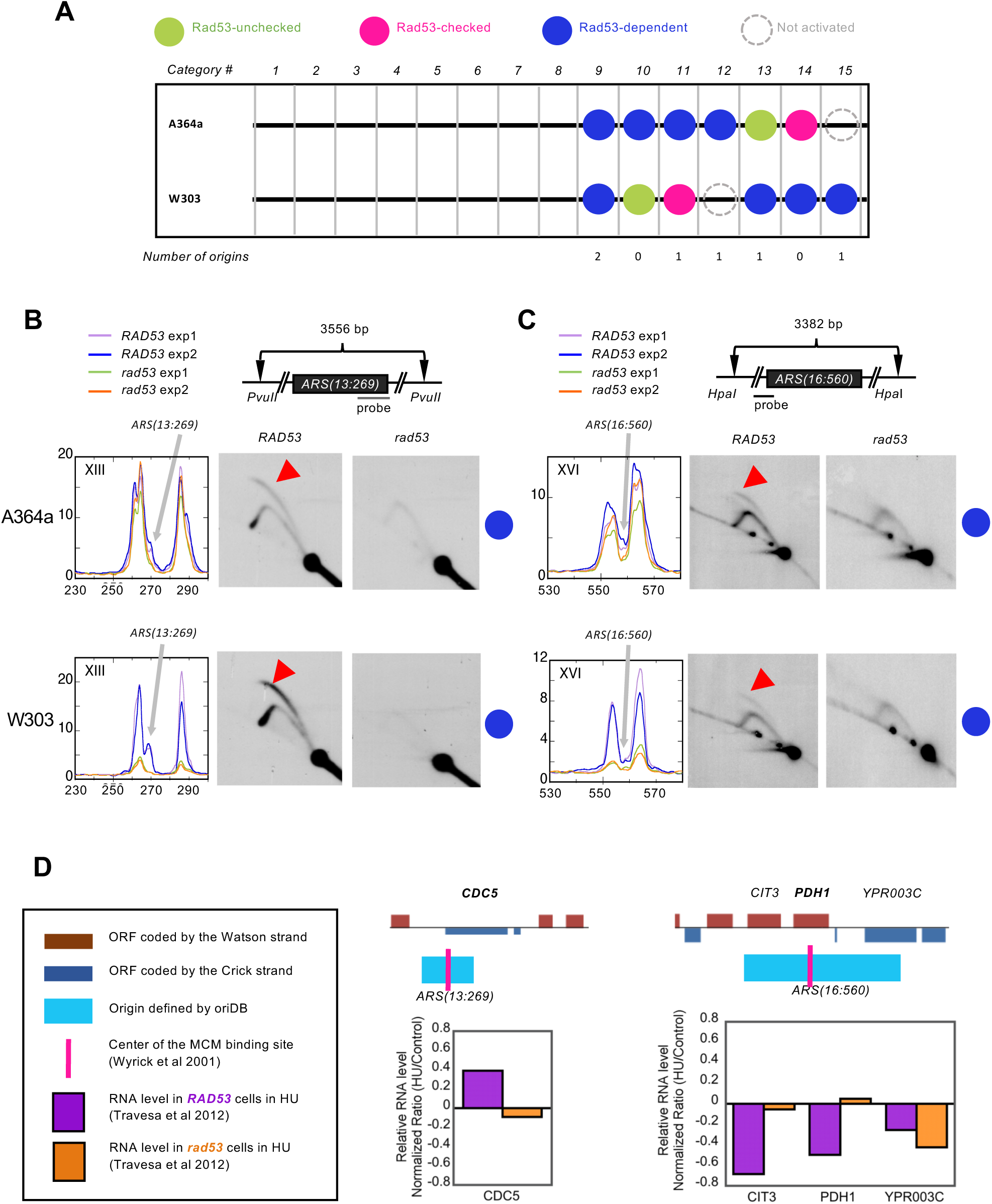
Rad53-dependent origins. **(A)** Numbers of Rad53-dependent origins parsed into the shown categories. (**B&C**) 2-D gel validation of *ARS(13:269)* and *ARS(16:560)* performed similarly as described in Fig. 3. **(D)** Gene expression levels at genomic loci of Rad53-dependent origins that overlap with ORFs. The data for gene expression are extracted from Travesa et al (31). The relative locations of the origins and location of ORC binding sites are derived from OriDB. The ORF that is nearest to the ORC binding site is bolded in each locus.

We next asked if the strain-specific usage could be explained by sequence polymorphisms within these origins. We made use of the previously published contig sequences from the *de novo* assembly of whole genome sequencing data in W303 [17]. To identify SNPs in the W303 strain we performed BLAST searches using DNA sequences containing 71 strain-specific origins retrieved from the S288c reference genome against the contigs from W303. We also generated contigs from *de novo* assembly of whole genome sequencing data of an A364a strain and performed BLAST searches similarly. We found 32 origins with full-length sequence contigs in both genetic background based on OriDB coordinates. Eight of these 32 origins contain strain-specific polymorphisms, with five (*ARS201.5, ARS(5:109), ARS810, ARS1233,* and *ARS(15:1023)*) in the A364a background and three (*ARS207.1, ARS221.5,* and *ARS1429*) in the W303 background (Fig. 4A&B). All but one origin, *ARS1429*, were W303-specific and showed no activity in A364a. Next, we searched for ACS elements in these origins and asked to what extent the strain-specific origin usage was due to polymorphisms in the ACSs. Unexpectedly, none of the eight origins contained any consensus ACS, despite having multiple near matches, in the OriDB-curated ARS coordinates. We then surveyed the ARS sequences from a high-resolution ARSseq study to obtain more precise locations for these eight origins and looked for consensus ACSs [33]. Unfortunately, only two origins (*ARS201.5* and *ARS1233*) were mapped by ARSseq. The average origin size based on ARSseq is significantly smaller than that based on OriDB (568 bp vs. 1274 bp), making it unlikely that we missed the consensus ACSs by using the OriDB coordinates. Indeed, the ARSseq core sequences for *ARS1233* were well within the boundaries defined by OriDB. *ARS201.5* ARSseq core sequences (28719-29138) extended beyond the region defined by OriDB (28941-29160) at the 5’-end. We searched the extended region and still found no consensus ACS. Therefore, it appeared that the strain-specific origins are devoid of consensus ACSs and thus are dependent on the near-matches of ACS elements to initiate replication. We then asked if the near-match ACSs were polymorphic between ARS364a and W303. Only *ARS201.5*, a W303-specific origin, contained a polymorphism (29064:T>A) within such a near-match ACS (AAAA<u>T</u>ATCAAG, ACS is on the - strand) in the A364a background (Fig. 4A). Note the 29064:T>A change actually improved the ACS element in A364a. None of the other polymorphisms occurred in a near-match ACS, nor did they create a new ACS. Taken together these data suggest that strain-specific origin usage cannot be accounted for by SNPs in the ACS elements.

Additionally, closer scrutiny of the ssDNA profiles revealed two distinct patterns of strain-specific usage of origins. In one of them a more binary origin activity was observed, *i.e.,* a ssDNA peak at the specific origin was present in W303 and absent in A364a (Fig. 4B). In the other pattern two origins, *ARS201.7* and *ARS1429*, which were inactive in A364a and W303, respectively, appeared to have shifted the activation site downstream from the origin. In both cases multiple SNPs were found in the genetic background where the origin was not active (Fig. 4A). For instance, *ARS1429* was deemed inactive in W303 by the algorithm —a well-defined ssDNA peak at *ARS1429* was only visible in one of the two experiments in W303—and consequently initiation took place from a likely origin (*ARS(14:773)*) in that experiment (Fig. 4A). This likely origin was not activated in A364a (Fig. 4A). Similarly, the ssDNA peak appeared to have shifted from *ARS201.5* to a downstream location in the A364a background. The new initiation site is located between the confirmed origins *ARS201.5* and *ARS201.7* (Fig. 4A). No ARS, including dubious ARSs in the OriDB, was found in this region. Therefore, it appeared that the polymorphisms at these two origins in the corresponding genetic background rendered initiation to take place elsewhere.

### Rad53-dependent origins

Finally, we discovered six origins that were only active in *RAD53* and not the checkpoint mutant cells in at least of one of the two strain background (Fig. 5A). We termed this novel class of origins “Rad53-dependent origins”. Two of these origins showed strain-specific usage, *ARS1016* (W303) and *ARS1618.5* (A364a), as discussed above (categories 12 and 15 in Fig. 5A). Another two origins, *ARS1206.5* and *ARS207.8*, showed Rad53-dependence in only one background (categories 11 and 13 in Fig. 5A). Finally, two origins were consistently identified as Rad53-dependent in both strain background: *ARS(13:269)* and *ARS(16:560)* (category 9 in Fig. 5A). We first focused on the two origins that were consistently identified as Rad53-dependent in both A364a and W303 and validated them by 2-D gel. Indeed, both *ARS(13:269)* and *ARS(16:560)* appeared to be only active in *RAD53*, and not in *rad53* cells (Fig. 5B&C). This is a new class of origins that have not been described in literature. The only discernable feature associated with these origins was their proximity to the nearest ORF. Five of these six Rad53-dependent origins, with the exception of *ARS1206.5*, overlap with an ORF. This led us to ask if the lack of origin activity in the *rad53* cells was due to higher level of transcription at these loci in *rad53* cells than in the *RAD53* cells using published data [34]. Very low expression level of *PDH1*, which is associated with the validated Rad53-dependet origin, *ARS(16:560),* was detected in either *RAD53* or *rad53* cells. The level of *CDC5*, the ORF that is associated with *ARS(13:269)*, was actually lower in *rad53* cells than in *RAD53* cells. For the remaining four origins the overlapping ORFs all showed decreased expression in *rad53* cells, compared to *RAD53* cells (Fig. 5D). These results suggested that the lack of origin activation in *rad53* cells is not a result of increased level of transcription.

## Discussion

We conducted a comparative analysis of origin activation, with and without DNA replications tress by HU, using genomic ssDNA mapping in two common laboratory yeast strains. By coupling with whole genome sequence information our experimental design offered the opportunity to directly test the impact of sequence polymorphism on origin usage as well as on the checkpoint response to replication stress. The ssDNA mapping results were corroborated by 2-D gel analysis. Consistent with previous findings more origins in the genome are under the control of the Rad53-mediated checkpoint in HU, *i.e.*, do not fire in HU in wild type *RAD53* cells, than origins that are impervious to the checkpoint in both genetic backgrounds. The vast majority of origins (331) share the same checkpoint status between the two genetic backgrounds. This suggests that the replication checkpoint by and large exerts control over the same origins in A364a vs. W303, provided the origin is competent for activation. Only 13 origins showed opposite checkpoint status in the two genetic backgrounds. Below we discuss the key observations with regards to the differences in genetic background.

We detected 71 origins that showed strain-specific usage, 32 of which were fully sequenced in both the W303 and A364a background. Comparative sequence analyses of 8 origins with strain-specific polymorphisms indicated that none of the strain-specific origin usage could be explained by polymorphisms in the consensus ACS elements. In fact, none of these 8 origins contained a canonical ACS, only near-matches of an ACS with 1-2 mismatches. This observation suggests that consensus ACSs are not essential for activation of these strain-specific origins. Instead, the presence of multiple near-match ACSs acting in concert is probably sufficient for origin activation. The vast majority of the SNPs, with only one exception, were located even outside these near-matches ACSs. Finally, the remaining 24 strain-specific origins are devoid of polymorphisms within the origin, suggesting that distal elements from the current known origin regions might play a role in conferring strain-specific usage. Therefore, our genome-scale analyses provided exceptions to the presumed essential role of the ACS for origin activation.

Compared to the A364a background there were more activated origins in W303, specifically in the Rad53-checked category. The W303 cells also showed a smaller distance covered by replication forks than A364a-*RAD53* cells. Because the ssDNA at replication forks is induced by HU, we directed our investigation at the Rad53-mediated checkpoint response. We detected a higher level of phosphorylated Rad53 in W303 cells than in A364a. Moreover, while Rad53 phosphorylation was greatly reduced by the checkpoint mutation in both W303 and A364a, the level of the mutant Rad53 protein was lower in W303. We speculate that these results could be explained by the dual functions of Rad53 in limiting origin activation and replication fork progression. The stronger activation of wild type Rad53 in W303 cells may cause increased inhibition of replication fork progression relative to the A364a background. Interestingly, a recent study reported that Rad53 limits excessive unwinding of DNA at replication forks by the replicative helicase CMG (Cdc45-MCM-GINS) complex [35]. On the other hand, the lower Rad53K227A protein level due to checkpoint mutation in W303 cells would cause more Rad53-checked origins to activate. It would be of interest to determine if titrating the Rad53 level can modulate the number of activated origins within the same strain.

Finally, we discovered a new class of “Rad53-dependent Origins”. The only recognizable feature of these origins is their proximity to an ORF. In the two validated Rad53-dependent origins in both A364a and W303, *ARS(13:269)* and *ARS(16:560)*, the MCM binding site is located inside *CDC5* and *PDH1*, respectively. Surprisingly, origin activity did not appear to correlate with low level of transcription, contrary to the idea that transcription interferes with origin activity. In fact, the gene expression level of *CDC5* was higher in *RAD53* cells than in *rad53* cells (Fig. 5D). Thus, it appears that transcription does not preclude the activities of these Rad53-dependent origins. Intragenic origins in the yeast genome are relatively less common, with *ARS604* and *ARS605* being the primary examples [29]. In both cases it was thought that transcription abolishes origin activity, constitutively at *ARS604* (inactive in both mitosis and meiosis) and meiotically at *ARS605* (only active during mitosis). However, this notion may require further scrutiny. First, *ARS604* activity has been detected in checkpoint mutants previously [21] and in the current study (File S1), calling into question the inactive status that has been associated with this origin. Moreover, MCM binding was detected at *ARS605* in meiosis [30] despite the conclusion that it is inactive during meiosis [36]. Therefore, we suggest that activation of intragenic origins is more common than previously thought, at least in the context of replication stress in mitotic cells. Nevertheless, we note that in almost every case of the “Rad53-dependent origins”, there is an active origin close-by. It is possible that the RAD53-dependent origins have a higher probability to fire because the nearby origin is more likely to be inactivated by transcription. In *rad53* cells, where transcription is reduced, the adjacent origin becomes more active, rendering the Rad53-dependent origin passively replicated. However, the underlying assumption for this explanation would be that low level transcription selectively impacts the adjacent origin, rather than the Rad53-dependent origin itself. For these reasons we believe that a more probable explanation for the Rad53-dependent nature of these origins is the requirement of an intact Rad53 checkpoint function. Whether Rad53 is required for establishing or maintaining the pre-Replication Complex assembly at these genomic locales remains to be determined.

In summary, our study provides a comprehensive catalogue of active origins in two common lab strains, which will assist future phylogenetic characterizations of replication origins in yeasts as well as our general understanding of the mechanisms of origin usage in eukaryotic organisms.

## Materials and Methods

### 1. Yeast strains

Yeast strains used in this study were derived from A139 (*MAT*a *RAD5 bar1::LEU2 can1-100 ade2-1::ADE2 his3-11,15 leu2-3,112 trp1-1::TRP1 ura3-1::URA3*) [37], a *RAD5* derivative of W303 [38] or HM14-3a (*MAT*a *bar1 trp1-289 leu2-3,112 his6*) in the A364a background [39]. The *rad53K227A* mutation was introduced into the *RAD53* gene locus by gene replacement as described [40].

### 2. Genomic ssDNA mapping by microarrays

Detailed procedures for ssDNA labeling, microarray hybridization and data analysis were described previously [32]. Briefly, cell cultures were grown to an OD_600_ of 0.25∼0.3 in YPD medium, followed by G1 arrest with 200 nM α-factor. Pronase (0.02 mg/mL) was used to synchronously release cells into S phase in the presence of 200 mM hydroxyurea (HU). G1 control and S phase samples were collected prior to cell cycle release and after 1 h treatment of HU, respectively. Three-hundred ml of cells from each sample were collected and spheroplasted in agarose plugs for ssDNA labeling. Differentially labeled G1 (Cy5-dUTP) and S phase (Cy3-dUTP) DNA were co-hybridized onto Agilent Yeast Whole Genome ChIP-to-chip 4 × 44K (G4493A) microarrays and the data were extracted by the Agilent Feature Extraction Software (v9.5.1). The relative quantity of ssDNA at a given genomic locus was calculated as the ratio of the fluorescent signal from the S phase sample to that of the G1 control, followed by Loess-smoothing over a 6-kb window at a step size of 250 bp.

### 3. Flow cytometry

Cells were grown identically as described above. One-ml of cells were collected from logarithmically grown cell culture (asynchronous control sample), and at 0, 15, 30, 60, 90, and 120 min following synchronous release from α-factor arrest into S phase in the presence of 200 mM HU. Cell fixation and processing for flow cytometry were performed as previously described [37].

### 4. Whole cell lysate preparation, SDS-PAGE and western blotting

Protein lysate preparation was adapted from Dr. Philip Zegerman’s protocol. For each sample/condition, 10^8^ cells were collected. Cells were pelleted by centrifugation at 13200 rpm for 5 min before quick freezing on dry ice. To prepare protein lysates, frozen cell pellets were thawed in 200 μl of 20% TCA and transferred to a rubber seal screw capped 1.5-ml Eppendorf tube. To each thawed sample, 400 μl of glass beads were added before vigorous agitation in a bead beater homogenizer in the cold room, twice for 30 sec each with a 45 sec rest between. The beads were then resuspended in 400 µl of 5% TCA and the supernatant was transferred to a fresh Eppendorf tube. The beads were then washed with 400 µl of 5% TCA, and the wash was pooled with the first supernatant. The pooled supernatant was centrifuged for 2 min at 13200 rpm and the pellet was resuspended in 200 µl of 1x Laemmli buffer containing β-mercaptoethanol and 50 µl of 1M Tris base (to increase the pH). Protein lysates were boiled at 98°C on a heat block for 10 min, cooled on ice, and centrifuged at 13200 rpm for 2 min to clear the lysate. Forty μL of cell lysate was loaded on an 7.5% SDS-PAGE gel and electrophoresed at 100 V until the 100 kDa marker reached the dye-front. Following transfer to 0.45-μm nitrocellulose membrane using standard procedure, the blots were blocked for 1 h at RT with 5% dry milk in 1x TBS-T, then hybridized overnight at 4°C first with 1:1000 Rabbit anti-Rad53 (Abcam #104232) or 1:4000 anti-Vma1 (gifted from the Kane lab), followed by 1:10000 anti-Rabbit (Invitrogen #31460) and 1:5000 anti-Mouse (Invitrogen #31430) for 1 h at RT, respectively. Blots were washed thrice with 5% TBS-T for 10 m each time. Post auto-radiography using ECL detection reagent (BioRad), the membranes were washed with dH_2_O, then incubated with Ponceau Stain Solution (0.1% Ponceau, 5% Acetic Acid) for 5-10 min before photographing.

### 5. Identification of significant ssDNA peaks

We first identified those ssDNA peaks with significant height in each sample by a Python module, PeakUtils 1.1.0 (https://pypi.python.org/pypi/PeakUtils). This module requires a user-input parameter, the minimal distance between two adjacent peaks. The size of this minimal distance is inversely correlated with the total number of ssDNA peaks identified, as fewer close-by peaks would be simultaneously called as peaks with increasing minimal distance. We determined the median inter-peak distance using a range of minimal distance set from 0.5 to 2.75 kb (Table S1) and compared to the inter-origin distance (∼27 kb and ∼15 kb for 410 confirmed origins and for 626 confirmed and likely origins, respectively). We then asked in each of the six groups how many ssDNA peaks were associated with the 626 origins, *i.e.*, the ssDNA peak summit was located within the origin region (Table S2, see below for detailed definition for origin regions). We chose a minimal distance of 1.75 kb between adjacent peaks as the most appropriate based on the following considerations. First, in this group the average inter-peak distance from two biological replicates of *rad53* cells (where we expect virtually all origins to fire) was 20.75 kb in both A364a and W303, which fell well within the expected range for inter-origin distance (15 to 27 kb as described above). Second, by increasing the minimal distance from 0.5 to 1.75 kb we significantly decreased ambiguous assignment of a ssDNA peak to multiple (>2) closely spaced origins while maintaining correct assignment of two “split peaks” to those known early/efficient origins with inferred bi-directional fork movement. For instance, using Experiment #1 for *rad53* cells in A364a as a training data set we found all 32 cases where a single origin was associated with 2 ssDNA peaks appeared to be due to bi-directional forks instead of close proximity to another origin (Fig. S4). However, further increasing the minimal inter-peak distance to 2.25 kb would result in false negatives as in the case of *ARS807*, because only one of the two split peaks would be associated with *ARS807*. Using the 1.75 kb minimal distance means those origins that are less than 1.75 kb away from each other would be potentially false negatives. Among the 626 confirmed and likely origins, 14 pairs of origins involving 26 unique origins have <1.75 kb inter-origin distance. Therefore, we estimated the false negative rate to be at lease 4% (26/626).

Another key determinant for ssDNA peak-origin association was the size of an origin. To avoid introducing potential bias due to differential origin sizes reported in OriDB, we defined each of the 626 origins as a region with a uniform size centering on the mid-point chromosome coordinate given by OriDB and we incrementally tested a range between 1 and 6 kb (Table S2). We chose 4 kb as the optimal size for two reasons. First, previous study has shown that the “replication bubble” size, or the overall distance that a pair of bi-directional replication forks would have covered in *RAD53* cells treated with 200 mM HU for 1 h, was ∼4.2 kb [23]. Second, setting origin size at 3 kb only identified 16 origins with bi-directional replication forks (compared to 32 at the 4 kb-origin size), based on the training data set of Experiment #1 for *rad53* cells in A364a (Table S2). In contrast, setting origin size at 5 kb enabled correct identification of 6 more origins (3 with bi-directional forks and 3 with single peaks), but also incurred 6 false positives (Table S2). Therefore, we chose the 4 kb origin size for further analysis.

The Lowess-smoothed ssDNA values from each experiment were input into the INDEXES function of the PeakUtils 1.1.0 module with a 1.75 kb minimal inter-peak distance. The amplitude index (0-1 in value) of a ssDNA peak was calculated as

(ssDNA - Min)/(Max -Min),

where Min and Max are the minimal and the maximal ssDNA value in the genome, respectively. A ssDNA peak amplitude index must exceed a threshold determined as follows for it to be considered significant:

Threshold = (Med + 3δ - Min)/(Max -Min), where

Med is the median ssDNA value of the given sample, and δ is the standard deviation of all ssDNA values below Med.

### 6. Origin identification by their association with significant ssDNA peaks

The mid-point locations of 626 confirmed and likely origins were obtained from OriDB (http://cerevisiae.oridb.org/index.php). All origins were then defined as a 4 kb region centering on the mid-point. A significant ssDNA peak was determined to be associated with an origin if its location had at least 1 bp overlap with the origin region, using the INTERSECT function in BEDtools [41]. An origin is considered active if it is associated with a significant ssDNA peak (defined in the previous section) in both biological replicate experiments.

### 7. Two-dimensional (2-D) agarose gel electrophoresis

Three hundred ml of cells were grown to an OD_600_ of 0.25 ∼ 0.3 in YPD medium and then synchronized in G1 with 200 nM α-factor. HU was added to a final concentration of 200 mM and then pronase was added at 0.02 mg/ml to release cells from α-factor arrest into the S phase. Cells were collected after 1 h. Genomic DNA was prepared according to one of two methods. In the first, cells were lysed and DNA purified according to the ‘NIB-n-grab’ method (http://fangman-brewer.genetics.washington.edu/nib-n-grab.html) and processed for 2-D gel analysis as described [11]. In the second method, cells from 100 ml of culture was collected and embedded in agarose before spheroplasts were prepared as described for ssDNA mapping. Each agarose plug was then washed with 5 ml of TE 0.1 (2X for 30 min each) followed by 5 ml of 1X restriction digestion buffer (2X for 1 h each) in a 6-well dish. The agarose plug was then transferred to a humidity chamber and incubated with 100 µl of restriction enzyme reaction mix at 37℃ overnight. An additional 30 µl of restriction enzyme reaction mix was added to each plug and incubated for 2-3 h. The plug was then washed with TE pH 8.0 and TE containing 0.1 mM EDTA, pH 8.0 for 1 h each before 2-D gel analysis as described [11].

DNA fragments containing *ARS305*, *ARS813, ARS1006*, and *ARS207.1, ARS(13:269)*, and *ARS(16:560)* were amplified by PCR from the yeast genome with the following primers and used for Southern blot:

ARS305-L: 5’-ATTCGCCTTTTGACAGGACG-3’;

ARS305-R: 5’-ATAACGGAGACTGGCGAACC-3’;

ARS813-L: 5’-GGGCAATTTACCACCTACGG-3’;

ARS813-R: 5’-ACGAAACTATTGGGGCCTCT-3’;

ARS1006-L: 5’-TCGGTTAATGAACACGTGGA-3’;

ARS1006-R: 5’-ATCCAACCAATGCCAACTGT-3’;

ARS207.1-L: 5’-GGCGGTAGCTCATTTTCTGA-3’;

ARS207.1-R: 5’-TTGTCCTGAAGGTGCTACCC-3’;

ARS(13:269)-L: 5’-CTTCGTTAAGGGCAAGACCA-3’;

ARS(13:269)-R: 5’-AGTTCTCCGATTGGCAGATG-3’

ARS(16:560)-L: 5’-CCGTCATGCCCCAAATACTG-3’;

ARS(16:560)-R: 5’-AGCCAACCATTCATCCCTCA-3’.

### 8. Whole genome sequencing and *de novo* assembly of the A364a genome

A364a *RAD53* cells (KK14-3a) were grown to mid-log phase in synthetic complete medium. Cells were harvested with the addition of 0.1% sodium azide and 50 mM EDTA. Cell were prepared for FACS by treating with 0.25 mg/ml RNase A for 1 h, 1 mg/ml proteinase K for 1 h, and stained with 1 μM SYTOX Green. G2 and S-phase cells were collected using a BD FACS Aria IIu machine. Genomic DNA was prepared using the Zymo Research YeaSTAR genomic DNA kit. Whole genome DNA libraries were prepared using the Illumina Nextera DNA Kit and sequenced at 2×50 bp paired-end on Illumina HiSeq 2500. Approximately 13 million reads for the S-phase sample were obtained and used for further analysis. *De novo* sequence assembly of the A364a genome was performed with Velvet [42] using default parameters.

### 9. Sequence analysis of strain-specific origins

Sequences for strain-specific origins, *i.e.*, origins that are activated in only one of the two genetic backgrounds (13 in A364a and 57 in W303), were obtained from the S288C reference genome R64.2.1 at the Saccharomyces Genome Database (https://downloads.yeastgenome.org/sequence/S288C_reference/genome_releases/). The range of sequences for each origin was determined based on OriDB classification. These origin sequences were then compared against the W303 contigs [17] and the A364a contigs (*de novo* assembly in this study) using the BLAST2.6.0+ engine at NCBI [43]. The top hit with sequence length difference less than 20 bp for each origin was used for further analysis. Origins with SNPs and InDels were thus identified through sequence alignment. ACSs in the 71 strain-specific origins were identified by FIMO scans from the MEME Suite [44].

### 10. Data extracted from a gene expression study by Travesa et al. [34]

Microarray-based gene expression data were extracted from the GEO database accession number GSE33695, samples GSM833009_hu.30vsut.0 (*RAD53* treated with HU for 30 min) and GSM833049_rad53.HU.30.1vsrad53.UT.0.1 (*rad53* treated with HU for 30 min). These samples most closely match the genetic background and experiment conditions in our study.

## Acknowledgements

We wish to thank M. K. Raghuraman and B. Brewer for helpful discussions and sharing the R script for heatmaps. We are also grateful to the Kane lab for the anti-Vma1 antibody and to B. Haarer for technical advice. We also thank P. Zegerman for sharing the protocol for immunoblotting for Rad53.

## Ethics approval

Not applicable.

## Accession numbers

Raw DNA sequences of the A364a genome and microarray raw data will be uploaded to the GEO database before publication.

